# Whole plant community transplants across climates reveal structural community stability due to large shifts in species assemblage

**DOI:** 10.1101/475046

**Authors:** Sara Tomiolo, Mark C. Bilton, Katja Tielbörger

**Affiliations:** Department of Ecology and Evolution, Plant Ecology Group, University of Tübingen, Auf der Morgenstelle 5, 72076 Tübingen, Germany; Department of Bioscience, Aarhus University, Vejlsøvej 25, 8600 Silkeborg, Denmark

**Author notes:** Corresponding author; Mark C. Bilton Katja Tielbörger.

**Keywords:** annual plant communities, climate change, Climatic Niche Groups, community ecology, reciprocal transplants, determinants of plant community diversity and assemblage, distribution range, community stability, field experiments

## Abstract

(1) Climate change will decrease precipitation and increase rainfall variability in Eastern Mediterranean regions, with responses of plant communities largely uncertain. Here, we tested short-term responses of dryland plant communities to contrasting rainfall regimes using a novel experimental approach.

(2) We exposed three annual plant communities to sharp changes in climatic conditions using whole community reciprocal transplants of soil and seed banks. We tested for the role of climate *vs.* community origin on community response and resistance. In parallel, we asked whether origin-specific climatic adaptations predict compositional shifts across climates.

(3) For both community origins, the most dry-adapted species in each community increased in dry climate and the wet-adapted species increased in wet climate. Dry community origins showed large compositional shifts while maintaining stable plant density, biomass and species richness across climates. Conversely, wet communities showed smaller compositional shifts, but larger variation in biomass and richness.

(4) Asynchrony in species abundances in response to rainfall variability could maintain structural community stability. This, in combination with seed dormancy, has the ability to delay extinction in response to climate change. However, increasing occurrence of extreme droughts may, in the long-term, lead to loss of wet-adapted species.

## Introduction

Understanding how climate may alter overall available plant resources (Sardans *et al*., 2008; Garcia *et al*., 2014) and impact upon community structure (Tilman & Downing, 1994; Gilman *et al*., 2010) is a major challenge in current ecological research (Maestre *et al*., 2012; Parmesan & Hanley, 2015). In cold regions for example, warming is likely to improve growing conditions and thus increase plant community biomass by increasing nutrient mobilization and expanding the length of the growing season (Garcia *et al*., 2014). In contrast, decreased rainfall in drier regions will likely have negative impacts on primary productivity, community composition, and their corresponding ecosystem services (Sala & Lauenroth, 1982; Peñuelas *et al*., 2007; Miranda *et al*., 2011). These effects may be particularly strong in those dryland ecosystems for which climate predictions indicate increasing incidents of droughts (Cubasch *et al*., 1996; Smiatek *et al*., 2011). Here, droughts and increasing temperatures will increase evapotranspiration, shorten the growing season and limit access to nutrients, thereby decreasing total community biomass (Peñuelas *et al*., 2007; Doblas-Miranda *et al*., 2015; Harrison *et al*., 2015). In extreme cases, this may lead to the collapse of entire ecological communities (Forey *et al*., 2010).

One of the predicted impacts of climate change is a re-assembly of plant communities (Hobbs *et al*., 2006; Williams & Jackson, 2007; Alexander *et al*., 2016) due to the differential ability of single species to either track their climatic niche or to survive under changed conditions, by means of adaptation or plasticity (Fernandez-Going *et al*., 2013; Shi *et al*., 2015). Such community reshuffling may be expressed in a loss or gain of certain species, a shift in species relative abundance or both. In plant communities already exposed to large inter-annual variations in climate, immediate rearrangement of community assembly in response to climate extremes may be an inherent property of plant communities, and may promote community stability in the long-term. This effect is particularly pronounced when species numbers are large and population sizes vary asynchronously (Doak *et al*., 1998; Schindler *et al*., 2015). Indeed, ecological theory and models support the idea that high inter-annual variation in species response to climate can lead to community-level stability (Anderson *et al*., 1982; Tilman *et al*., 1998; Thompson *et al*., 2015; Abbott *et al*., 2017). This may be an important mechanism for maintaining dryland communities’ stability in response to large year-to-year variation in rainfall, and for slowing down ongoing selection processes due to climate change (Bonebrake & Mastrandrea, 2010; Bilton *et al*., 2016).

Although long-term climate manipulations are the gold standard in ecological climate impact research and are fundamental for understanding long-term community shifts (Brown *et al*., 2001; Rinnan *et al*., 2007; Blume-Werry *et al*., 2016), they are very costly to set up and maintain, often outliving funding cycles and scientific research positions (Lindenmayer *et al*., 2012). The monitoring of communities for short-term responses may be a useful complement to long-term experiments, as besides being less costly, it can be vitally important for parsing mechanistic information about plant responses to large inter-annual variation, as well as extreme events (De Dato *et al*., 2006; Barbosa *et al*., 2014; Blume-Werry *et al*., 2016). Reciprocal transplants represent a promising approach for indirectly studying plant responses to climate change on a short temporal scale. These manipulative experiments have been widely adopted in single species (e.g. Link *et al*., 2003; Casper & Castelli, 2007; Macel *et al*., 2007; Alexander *et al*., 2015; Tomiolo *et al*., 2015) for studying local adaptation and, more recently, for studying their responses to a climate that matches conditions predicted by climate change scenarios (the so called “space-for-time approach”). Reciprocal transplants have also been applied to entire communities in studies of soil microbiomes (Waldrop & Firestone, 2006; Lazzaro *et al*., 2011), leaf litter (Ayres *et al*., 2009; Allison *et al*., 2013), and occasionally to whole plant communities in different habitats ranging from wetlands to alpine grasslands (Maranon & Bartolome, 1993; Wetzel *et al*., 2004; Wu *et al*., 2012; Alexander *et al*., 2015). However, the potential for using whole community reciprocal transplants to study plant community response to climate change has not been fully exploited, particularly in dryland systems, which often provide ideal conditions.

Dryland ecosystems are often dominated by annual plants that survive the dry season as a permanent seed bank (Cohen, 1966). Therefore, the community (i.e. the seed bank) can be conveniently transplanted as a whole during the dry season without any damage to the plants. In addition, by transplanting seed banks with their associated soil, it is possible to evaluate plant communities’ responses to climate while preserving soil abiotic and biotic interactions. To test the response of dryland annual plant communities characterized by very different climates, we transplanted home soil with seed bank among three sites situated along a steep aridity gradient in the Eastern Mediterranean region, ranging from arid to Mediterranean climate. In this region, rainfall is the main limiting factor for plant growth (Ziv *et al*., 2014) and differs up to eight-fold between the driest and wettest site (Holzapfel *et al*., 2006). We classified species based on their climatic requirements, adopting the Climatic Niche Group approach (CNG; *sensu* Bilton et al. 2016) that has been successfully employed for the species in our study region (Bilton *et al*., 2016) and in other dryland ecosystems (Liu *et al*., 2018). By identifying those species responsive to drier or wetter conditions, this approach provided us with testable predictions about directions of shifts in community assembly across climates within the reciprocal transplants. Finally, the study sites used for our reciprocal transplant also hosted a long-term climate manipulation experiment (Tielbörger *et al*., 2014). This allowed for a qualitative comparison between long-term dynamics, resulting from consistently imposed climate change, and the short-term responses observed in our transplant experiment.

We predicted that the community emerging from the reciprocal soil transplants would be greatly determined by community origin, with fewer individuals emerging from drier origins than wetter origins. Secondly, we hypothesized that, regardless of their origin, communities emerging at the drier transplant site (i.e. lower rainfall availability) would experience a reduction in total biomass and plant density. We also predicted that climate would select the emerging community from the species pool of each origin in a predictable manner, with more wet adapted species emerging when communities were exposed to wetter climates, and more dry-adapted species in drier climates.

## Methods

### Study area

This study was conducted in Israel at three fenced sites (area approximately 100 m × 400 m) with respectively Mediterranean (M), semi-arid (SA) and arid (A) climate. The three study sites share the same calcareous bedrock, southern aspect, altitude and mean annual temperatures, so that they differ chiefly in mean and variance of annual rainfall, and vegetation. The M site is located southwest of Jerusalem (N 31° 42’ E 35° 3’) at 620 masl, on Terra Rossa soil. The climate is characterized by 550 mm average annual rainfall with 20% inter-annual variation. The SA site (N 31° 23’ E 34° 54’) is located in the northern portion of the Negev Desert near the city of Beersheba, at 590 masl, on Light Brown Rendzina. Average annual rainfall is 270 mm with approximately 30% inter-annual variation. The A site is located in the central Negev near Sde Boqer (N 30°52’ E 34°46’) at 470 masl, on desert Lithosol. Average annual rainfall amounts to 90 mm with 43% inter-annual variation (Holzapfel *et al*., 2006). The plant communities at the three sites are semi-natural shrublands dominated by *Sarcopoterium spinosum* (L.) Spach, and winter annuals (approx. 85% of all species) that persist during summer in the form of dormant seed banks stored in the soil (Noy-Meir, 1973; Alon & Steinberger, 1999). The species pool is overlapping among sites, and annual plant cover amounts to 25% at the M site, 10% at the SA site, and < 1% at the A site (Tielbörger *et al*., 2014).

### Experimental set up

During summer of 2010, we collected soil with seed bank from forty square plots (20cm × 20cm, depth: 5cm) at the M and A sites and sixty plots at the SA site. Within each site, plots were situated at least 20 cm apart from each other, and away from rocks and shrubs. Following Tomiolo et al. (2015) soil collected from each site was pooled to produce a baseline community. Previous studies showed that small-scale heterogeneity in the soil seed bank may be very large, with some patches having almost no seeds and others very many (Siewert & Tielbörger, 2010). Therefore, we pooled the soil samples per site prior to the transplant, following the procedure adopted in many previous studies using field soil (Maranon & Bartolome, 1993; Macel *et al*., 2007; Burns & Strauss, 2011; Lazzaro *et al*., 2011).

The soil was stored in a net-house at the University of Rehovot, Israel, where it experienced summer temperatures necessary for breaking seed dormancy (Baskin *et al*., 1993). In September 2010, twenty of the previously excavated plots at each site were randomly selected and filled with home soil, while the remaining plots were filled with soil from the closest away-from-home site (i.e. M site received M and SA soil; SA site received M, SA and A soil; A site received SA and A soil, Supplementary Material Appendix 1 Fig. A1). Transplanted soil was separated from the surrounding soil by a layer of absorbent paper that provided initial isolation between soils, while not impeding water percolation. After transplanting, we placed patches of organza (a thin transparent fabric) over the surface of each plot to avoid contamination from seed dispersal or seed predation (Petrů & Tielbörger, 2008), and we removed them at the time of germination.

Because the transplants were carried out during the dormant season we could relocate the community of dormant seeds and soil biota with minimum damage. By transplanting communities with their maternal soil we could test direct effects of climate (e.g. decreasing rainfall) while preserving biotic interactions with neighbouring plants and soil biota, which are also affected by the novel climate (Emmett *et al*., 2004). At peak development (spring 2011), we recorded the identity and number of individuals of the emerging species in each plot. In order to minimize edge effects, we excluded plants growing in the outer 1 cm margin of each plot. After species identification, aboveground biomass was collected, oven-dried at 70°C for 48 hours and weighed.

Unfortunately, the season of recording was very dry and the arid site received only 30% of the average annual rainfall. Therefore, only a handful of seedlings of two desert species (*Stipa capensis*, *Erodium touchyanum*) emerged at the arid site. As a result, there was no home arid community to be compared to the transplants, and we had to restrict our subsequent analyses to the reciprocal transplants between the SA and M community origins.

### Climatic Niche Groups (CNG)

Each species within the target communities was assigned to a Climatic Niche Group (Bilton *et al*., 2016) classified by their distribution range in relation to rainfall. This approach has proven powerful for predicting species-specific response to climate change (Bilton *et al*., 2016; Liu *et al*., 2018). A similar method has been employed for defining thermal niches of species in high elevation and tundra habitats (Gottfried *et al*., 2012; Elmendorf *et al*., 2015), and it is conceptually similar to Ellenberg values, which determine species habitat requirements based on several abiotic parameters (Ellenberg, 1974).

The rationale for the CNG grouping is that rainfall is the main driver of community composition in the region, therefore species sharing similar climate adaptations (approximated by the realized climatic rainfall niche) are likely to co-occur in the same community (García-Camacho *et al*., 2017). Species realized climatic niche values were derived as in Bilton *et al*. (2016). For each single species the observed occurrences within Israel (distribution range) were overlaid with mean annual rainfall climate data, and the mean value was taken (obtained from BioGIS, 2012, available at http://www.biogis.huji.ac.il/). Boundaries between climatic niche groups spanned similar ranges of average annual rainfall (approximately 130 mm) and resulted in four groups that ranked species with respect to their hypothesized response to climate. Climatic Niche Group 1 (CNG1) represented species associated with the lowest rainfall extremes of the gradient, conversely CNG4 gathered species distributed in areas with high rainfall. Species from all four CNGs were present in both communities (Supplementary Material, Appendix 2 Table A1), but varied in their proportional representation at each site, and could therefore be compared across sites and climates (Bilton *et al*., 2016).

### Statistical analyses

We first analyzed how total density (number of individuals per plot), total biomass, species richness (number of species per plot) and diversity (Shannon-Wiener Index) varied in response to climate, community origin and their interaction. In addition, we analyzed how the number of individuals belonging to each climatic niche group per plot (i.e. CNG density) varied in response to climate, community origin, with respect to the four-level categorical explanatory variable CNG identity (i.e. CNG1 – CNG4), including all two-way and three-way interactions. We applied generalized linear models with negative binomial distribution to total individual, CNG density and species richness using the MASS package (Venables *et al*., 2002) within the R software version 3.3.3 (R Development Core Team, 2014). Biomass and species diversity were analyzed using linear models. To meet model requirements biomass square root transformed. The significance of the models was assessed with a Type 3 ANOVA, using the “car” package (Fox & Weisberg, 2011).

Visual representation of the CNG density interactions was done using log-ratios calculated from the overall mean abundance of each group in each climate or origin. Showing relative change in overall abundances was also extremely helpful for visualizing the significant interactions we found with our models.

For testing how species composition varied with community origin and climate we used Redundancy Analysis (RDA, (Legendre *et al*., 2011)) in the R package ‘vegan’ (Oksanen *et al*., 2015). The interaction term was included in a full model and confirmed using a step-wise approach. The data were Hellinger transformed (Legendre & Gallagher, 2001) and scaled within plots. Significance of the model was tested using 999 permutations. To test if species composition could be explained by rainfall distribution range we regressed the resulting RDA ‘species mean scores’ against the ‘climatic niche value’ of each species, both for individual species and for the CNG classifications. Furthermore, we performed an RDA on the community-weighted means (Garnier *et al*., 2007) using the species ‘climatic niche value’ as a pseudo-trait.

## Results

Overall, 97 species were recorded in our transplant plots, among which 12.3% were grasses, 23.7% legumes and 64% belonged to other families (Supplementary Material, Appendix 2 Table A1). In total, 68 species emerged from the semi-arid soil seed bank, 81 from the Mediterranean origin, and 53 species were shared between the two origins. Fourteen of these appeared in all four combinations of community origin and climate.

### Total plant density, diversity, richness and biomass

Number of species, species diversity, total plant density and total biomass (Fig. 1, Table 1) were all significantly higher for M community origins rather than SA origins. Additionally, M community origins attained significantly lower species richness and diversity and biomass (Fig. 1) when exposed to the drier SA climate compared to their home M climate. For the SA community origins, climate had no significant effect on total plant density, biomass, species richness or diversity (Fig.1, Table 1).

**Figure 1:**
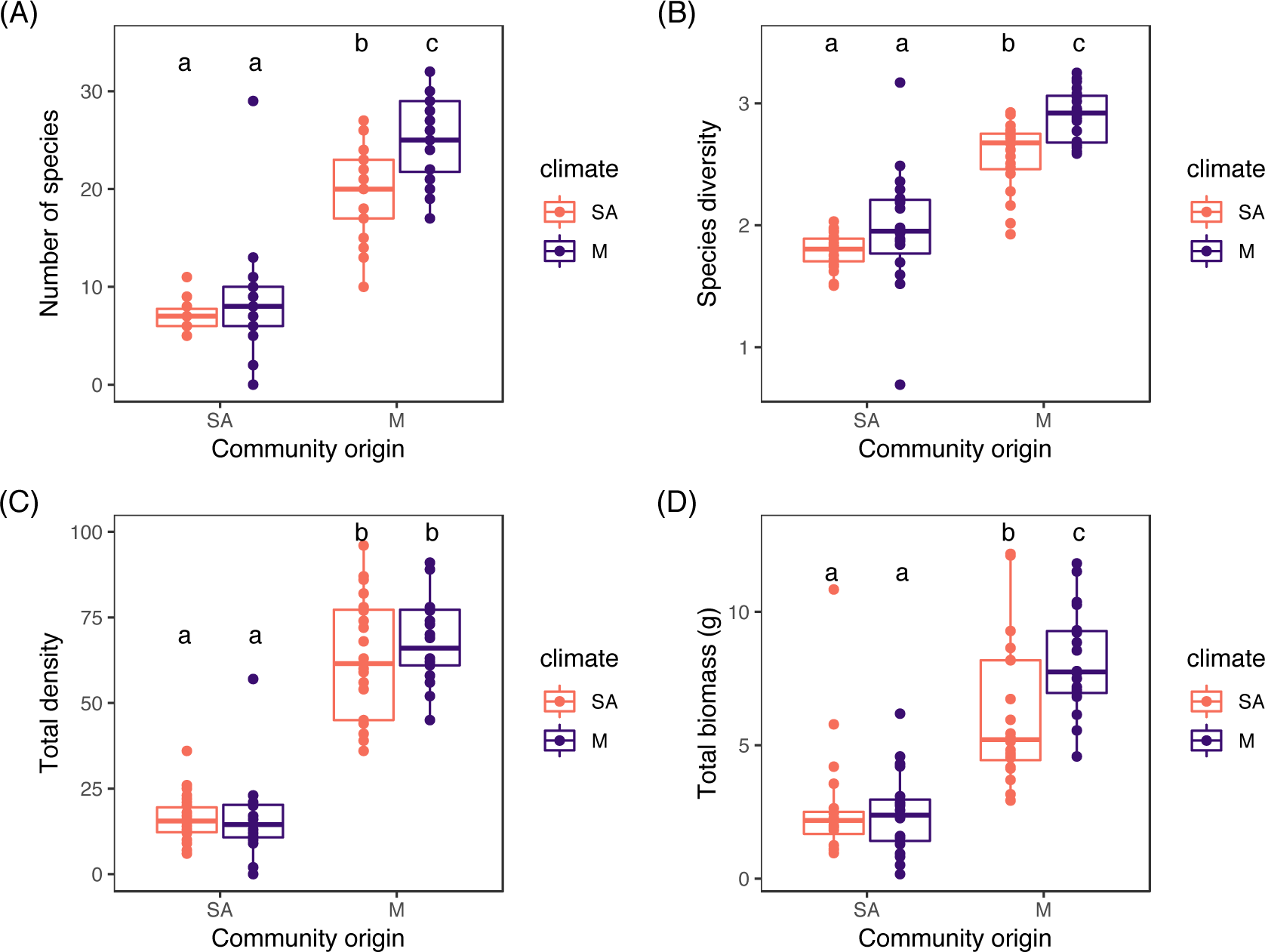
Mean ± 1SE A) species richness (i.e. number of species per plot), B) diversity C) total individual density, D) total biomass of annual plant communities established from two community origins (‘M’ Mediterranean; ‘SA’ Semi-Arid) grown in two sites (i.e. climates: ‘M’ Mediterranean; ‘SA’ Semi-Arid).

**Table 1:** Type III ANOVA table results for the models applied to individual density, species richness, species diversity, and total biomass. Lines correspond to response variables and columns to explanatory variables of each model. In each column the first value represent Chi-square test values and the second the p-value. Probability values for significant terms are reported in bold.

|  | Origin | Climate | Origin x Climate |
| --- | --- | --- | --- |
| Density | 312.86; <**0.001** | 0.12; 0.72 | 0.59; 0.44 |
| Species richness | 281.29; <**0.001** | 12.99; <**0.001** | 0.085; 0.77 |
| Species diversity | 130.41; <**0.001** | 10.40; **0.001** | 1.36; 0.24 |
| Total biomass | 103.86; **0.001** | 2.34; 0.13 | 4.66; **0.03** |

### Species composition

The RDA indicated four distinct communities emerging from the respective treatments, with a significant effect of community origin and climate on species assembly, as well as a significant interaction between these terms (Fig. 2a, b). Using simple correlations we assessed which plots/species scores changed and had most impact on each axis. We obtained three main RDA axes describing the species composition. For plot mean scores, RDA1 (9.2% explained) was correlated to overall differences between community origins, whereas the constrained RDA2 (3.3%; explained) and RDA3 (1.7%; explained) distinguished the climate × community origin interaction term. For species mean scores, RDA1 was positively correlated to species Climatic Niche values, and the correlation was positive but less strong for RDA2 and RDA3 (Fig. 2c). Results were further validated by an RDA on the community weighted mean traits using species Climatic Niche values as a trait, and showed significant community origin and climate effects (p<0.05). In combination, these results suggest that rainfall niche partially explained variation in species composition across treatments.

**Figure 2:**
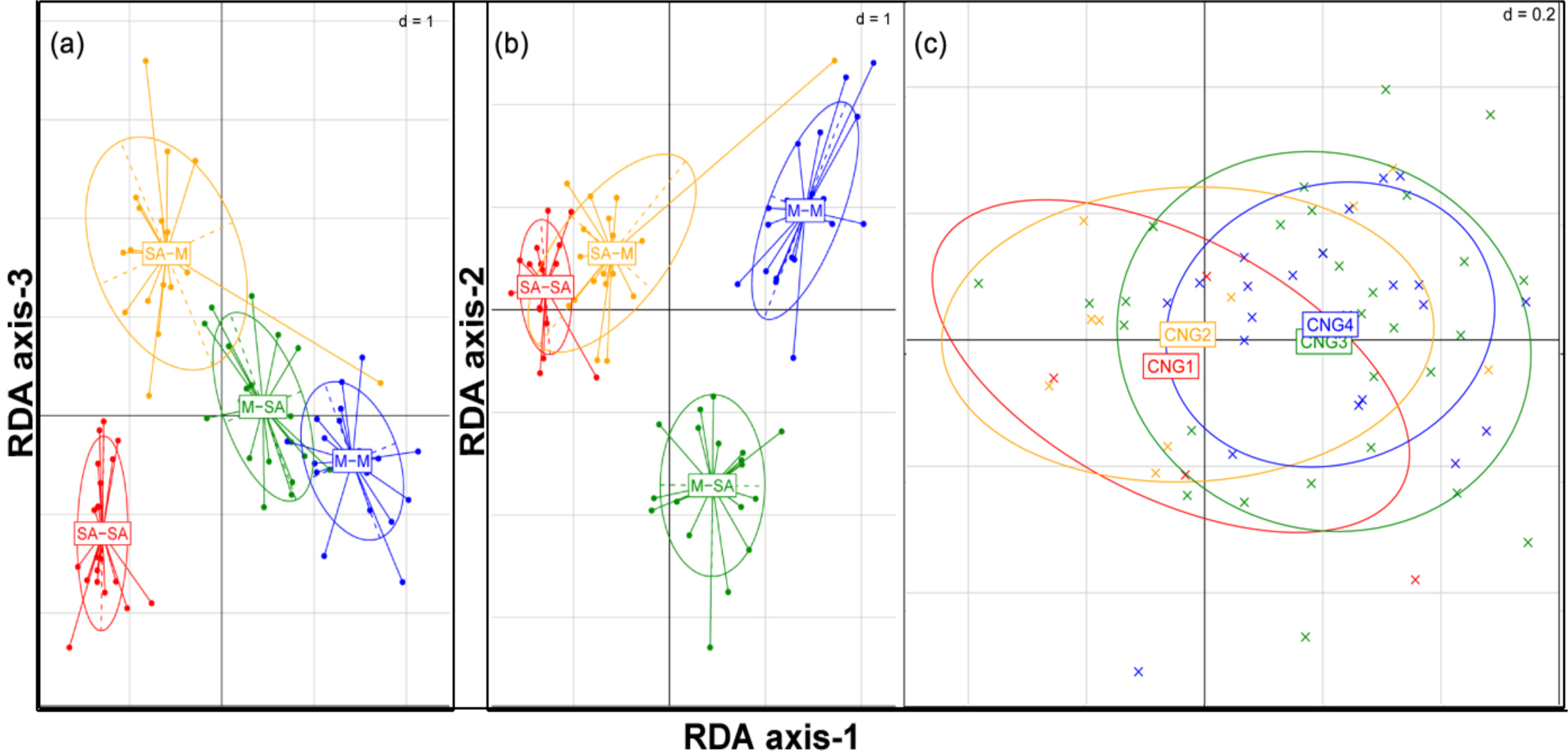
Graphical representation of Redundancy Analysis (RDA) showing species composition in community origins (M, SA) exposed to two different climates (M, SA). The ellipses represent the 95% confidence intervals for the groupings/categories. Fig. 2 a, b represents the plot centroids of each community origin-climate combination. Lines are vectors from the centre of a category to each site score (points). RDA-axis 1: correlated to distance between origins, RDA-axis 3: the effect of climate on SA community origins, and RDA-axis 2 the effect of climate on M community origins. In red: SA community origins - SA climate; yellow: SA origins - M climate; blue: M origins- M climate; green: M origins -SA climate. Fig. 2c represents the species centroids for each Climatic Niche Group (CNG). Lines are vectors connecting the centre of each group with species scores. In red: CNG1, yellow: CNG2, green: CNG3, blue: CNG4.

### CNG density across sites and community origins

Overall, densities of individuals in each CNG group significantly differed (CNG identity effect: Fig. 3a, b, Table 2), and all group densities were higher in M origin than SA origin (Origin and Origin × CNG effect; Fig. 3c; Table 2). The representation of CNGs in the communities also changed significantly across climates, dry groups were more abundant in SA climate and wet groups were more abundant in M climate (Climate × CNG effect: Fig. 3c, Table 2, Supplementary Material Appendix 2 Table A2). The magnitude in CNG shifts across climates was different among community origins as indicated by a significant 3-way interaction (CNG identity × community origin × climate, Table 2, Fig. 3e, f). In SA community origins, the mean abundance of individuals belonging to dry CNGs (CNG 1 and 2) was halved in the wet (M) compared to the dry (SA) climate; on the other hand individuals belonging to CNG 4, the wettest adapted group, were 6.5 times more abundant in SA communities origins emerging in the wet climate (Fig. 3a, e). In M community origins, the shift in CNG densities across climates was less strong compared to SA origins, but the hierarchical response of the CNGs was in the same order (i.e. densities of dry CNGs were higher in the dry climate and densities of wet CNGs were higher in the wet climate). The largest shift in density was seen in the wettest CNG that counted twice as many individuals in wet *vs.* dry climate (Fig. 3b, f).

**Figure 3:**
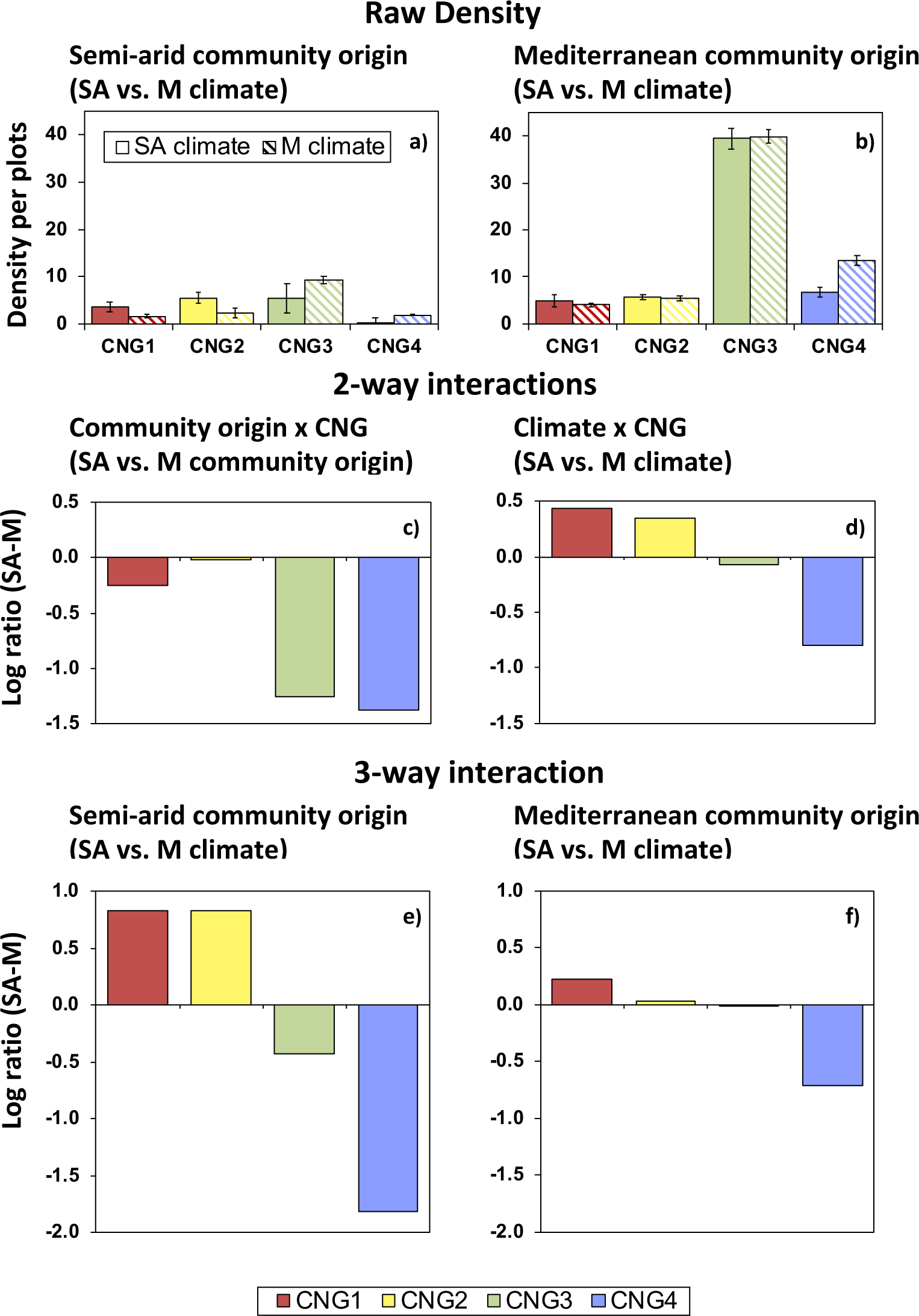
Impact of community origin, climate and Climatic Niche Group (CNG) on plant densities establishing from seed banks in home *vs*. away-from-home climate. Each Climatic Niche Group (CNG) aggregates species with similar climatic adaptation, ranging from dry climates (CNG 1) to wetter climates (CNG 4). Fig. 3 a, b: Total mean individuals’ densities across climates and origins are broken down according to CNG abundances. Fig. 3 c-f: Shifts in the abundance of CNG densities across community origins (3c), climates (3d) and the combination of the two (3 e, f). Shifts in densities of CNG groups are expressed as log ratios, thus visualizing changes in plant density on a relative scale. Positive values indicate higher CNG abundances in SA community origins or climate, whereas negative values indicate higher CNG abundances in M community origins or climate.

**Table 2:** Type III ANOVA table of results for Generalized Linear Models determining the effect of community origin, climate and Climatic Niche Group (CNG) on plant community densities establishing from seed bank.

| Effect | DF | LR Chi-sq. | P-value |
| --- | --- | --- | --- |
|  |  | 236.9 |  |
| Origin | 1 | 3 | **<0.001** |
| Climate | 1 | 2.35 | 0.12540 |
|  |  | 343.2 |  |
| CNG | 3 | 1 | **<0.001** |
| Origin x Climate | 1 | 0.07 | 0.78714 |
| Origin x CNG | 3 | 83.55 | **<0.001** |
| Climate x CNG | 3 | 50.09 | **<0.001** |
| Origin x Climate x CNG | 3 | 16.85 | **<0.001** |

## Discussion

Our results revealed that climate played a large role in determining the species assemblages in our whole community transplant experiment, and that such changes were predicted by species-specific climatic adaptations. One of our most notable findings was that the communities showed different qualitative and quantitative responses to climate change depending on their origin. Mediterranean community origins from a wetter and more predictable climate, responded with changes in species richness, diversity and total biomass, but showed small shifts in community composition. Conversely, semi-arid community origins, from a drier and more unpredictable climate, showed little variation in species richness, diversity and biomass, and large shifts in species and CNG composition.

Perhaps unsurprisingly, and as predicted from previous observations across the rainfall gradient (Tielbörger *et al*., 2014), communities establishing from the Mediterranean (M) origins had higher individual density and total biomass than semi-arid (SA) origins. This is also consistent with other dryland systems (Guo & Brown, 1997; Cleland *et al*., 2013). The higher plant density in Mediterranean community origins corresponded with higher densities of individuals for all CNG groups, Interestingly, the species composition and CNG abundances in the experimental communities rapidly matched the concomitant climate. Namely, plants establishing from both community origins showed a relative increase of dry adapted species (CNG 1 and 2) when exposed to the drier (SA) climate. Similarly, wet adapted species (CNG 3 and 4) were more abundant in both communities when exposed to the wetter (M) climate. This response was particularly large for semi-arid community origins where the reshuffling in community composition was strikingly well explained by hierarchical switches in CNG abundances. Therefore, CNGs revealed species responses to short-term climate effects in a predictable way, suggesting potential short-term selection mechanisms (e.g. environmental filtering) that act on the communities in response to yearly differences in rainfall. This result is consistent with the findings of Bilton *et al*. (2016), and suggests that CNGs are representative of species rainfall requirements and possibly climatic adaptations at the different sites.

Our most intriguing finding was that climate filtered for predictable species groups, but the magnitude of the structural community shifts was largely different among community origins. In addition, the magnitude of the compositional shifts between transplanted community origins was inversely related to changes in total community parameters across climates. Mediterranean origin transplants, had smaller compositional shifts, but larger shifts in biomass and richness. Conversely, semi-arid community origins, with higher between- and within-year rainfall variability (Tielbörger *et al*., 2014), experienced the greatest shifts in community composition, while showing only marginal response in community parameters. Consistent with our results, large interannual compositional shifts were observed also in other communities from drier and highly variable climates (Guo & Brown, 1997; Cleland *et al*., 2013). Interestingly, we did not record an increase in total biomass or total density in semi-arid community origins when exposed to wetter climates. This is possibly the result of an increase in wet CNGs in the communities, which have a core distribution in wetter regions where there are larger plant densities, productivity, and a higher intensity of competition (Schiffers & Tielbörger, 2006; Liancourt & Tielbörger, 2009). Wet CNGs may possess a better competitive ability and might have curbed the growth and density of dry CNGs, thus leading to little changes in total biomass and total density.

Large inter-annual variation in species abundances in the short-term may lead to higher community stability in the long-term (Bai *et al*., 2004; Grime *et al*., 2008). Similar patterns have been previously explained in plant community studies, albeit in a different context, by the portfolio effect (Doak *et al*., 1998; Schindler *et al*., 2015). The portfolio effect predicts that greater numbers of species in a community lead both mathematically and ecologically to a greater chance of asynchronous relationships forming year-to-year. Here we show, consistent with previous studies (Cleland *et al*., 2013; Hallett *et al*., 2014), that in the community originating in a more unpredictable climate (in our case the semi-arid community origin), greater asynchrony and greater species turnover led to greater stability across climates. Plant species in more variable climates have been found to exhibit a larger phenotypic plasticity (Sultan, 1987; Pratt & Mooney, 2013; Lazaro-Nogal *et al*., 2015; Spence *et al*., 2016), which results in fitness homeostasis (Richards *et al*., 2006; Nicotra *et al*., 2010). This could be an explanation for the higher resistance of species in SA communities but also reveals an interesting analogy with the community level, where ‘homeostasis’ may be associated with a larger compositional change. Overall, we suggest that asynchronous shifts in abundance of species according to their climate adaptations may allow for fast responses to year-to-year climatic variation in dryland annual communities (Abbott *et al*., 2017).

In the short-term, high species turnover may assure community stability in biomass and density, but in the long-term, such processes may also lead to greater resistance and stability of dry communities to rainfall fluctuations by favoring species adapted to arid conditions. This high turnover is possible without immediate loss of species because in dryland environments, plants often display bet-hedging strategies such as long-lived seed banks and seed dormancy that can buffer against inter-annual fluctuations (Petrů & Tielbörger, 2008; Tielbörger *et al*., 2012). Delayed germination of dormant seeds during unfavorable years decreases the risk of extinction over time and also promotes coexistence of species with different climatic requirements via storage effects (Chesson & Grubb, 1990; Pake & Venable, 1995). Interestingly, the findings from this short-term community transplant study are consistent with those of a parallel long-term experiment conducted at the same study sites (Tielbörger *et al*., 2014), where community composition was monitored for 10 years in permanent plots receiving respectively ambient rainfall, experimental drought (−30% rainfall) or increased rainfall (+30% rainfall). Plant communities exposed to the long-term climate manipulation treatments showed no detectable long-term effect on total density, species richness and community biomass (Tielbörger *et al*., 2014). However, at both time-scales, shifts in species abundance in relation to their CNG was observed (Bilton *et al*., 2016). While in our short-term study we found these effects to be more pronounced for semi-arid community origins, the long-term study found stronger patterns in Mediterranean communities (Bilton *et al*., 2016). This suggests that high inter-annual community fluctuations may contribute to stability in community composition in the long run, whereas low species turnover across years may result in long-term loss of wet-adapted species.

Our overall findings allow some careful conclusions about the potential response of these communities to climate change. It should be noted that the variation in rainfall experienced by the community origins in our study approximated the extremes of climatic variability at each site, but exceeded the decrease in rainfall predicted by climate change scenarios for the next 50-80 years (Smiatek *et al*., 2011; Tomiolo *et al*., 2015). These results suggest that, as long as inter-annual climatic fluctuations keep within the limits of climatic variability commonly experienced by these communities, and rainy years that replenish the seed bank periodically occur, wet adapted species will persist within the communities. However, with increasing drought and unpredictability, communities are likely to experience species loss that will affect primarily species with high rainfall requirements (Tielbörger *et al*., 2014; Bilton *et al*., 2016). The similarity of results between the long-term experiment and our reciprocal transplant indicate that the latter may be a powerful complement to long-term field experiments. However, annual communities are particularly suited for our experimental approach and the same may not hold for long-lived communities. For example, in temperate systems a lag between shifts in climatic conditions and subsequent changes in community structure is often observed (Adler & Levine, 2007; Jones *et al*., 2016). The unexpected community resistance and resilience of dryland ecosystems to extreme events compared with temperate ecosystems (Ruppert *et al*., 2015), may be attributable to the large variability in rainfall to which plant species are pre-adapted via bet hedging strategies (Tielbörger *et al*., 2012; Gremer & Venable, 2014) or enhanced phenotypic plasticity (Petrů *et al*., 2006). Such differences among rainfall variability and plant life history should be taken into account when drawing comparisons among habitats.

## Supporting information

Scheme of the reciprocal community transplant

Species list and mean number of individuals among Climatic Niche Groups

## Declarations

KT and ST developed the experimental design. ST set up the experiments and collected the data. MB and ST performed statistical analyses. ST wrote the first draft of the manuscript and all authors contributed substantially to the following versions. We declare that we do not have conflicts of interest.

## Acknowledgements

Jaime Kigel and Marcelo Sternberg provided logistic support. The Hebrew University of Jerusalem (Rehovot) kindly provided material for fieldwork. We thank Jake Alexander for providing comments on a previous version of the manuscript. This study is part of the GLOWA Jordan River Project and was funded by the German Ministry of Education and Research (BMBF). Further support for MB and ST was obtained by the German Research Foundation (TI338_12-1; TI338_11-1; and TI338_11-2; TI 338/15-1).

## Data Accessibility

upon publication data will be made accessible on Dryad

**Figure A1:** Dots on the map represent the three experimental sites. On the right hand, the the scheme of the community transplants at each sites. Communities (soil and relative seed bank) were transplanted between the home and the closest away-from-home site.

