## Supplementary figures and images for "Whole plant community transplants across climates reveal structural community stability due to large shifts in species assemblage"

### Scheme of the reciprocal community transplant

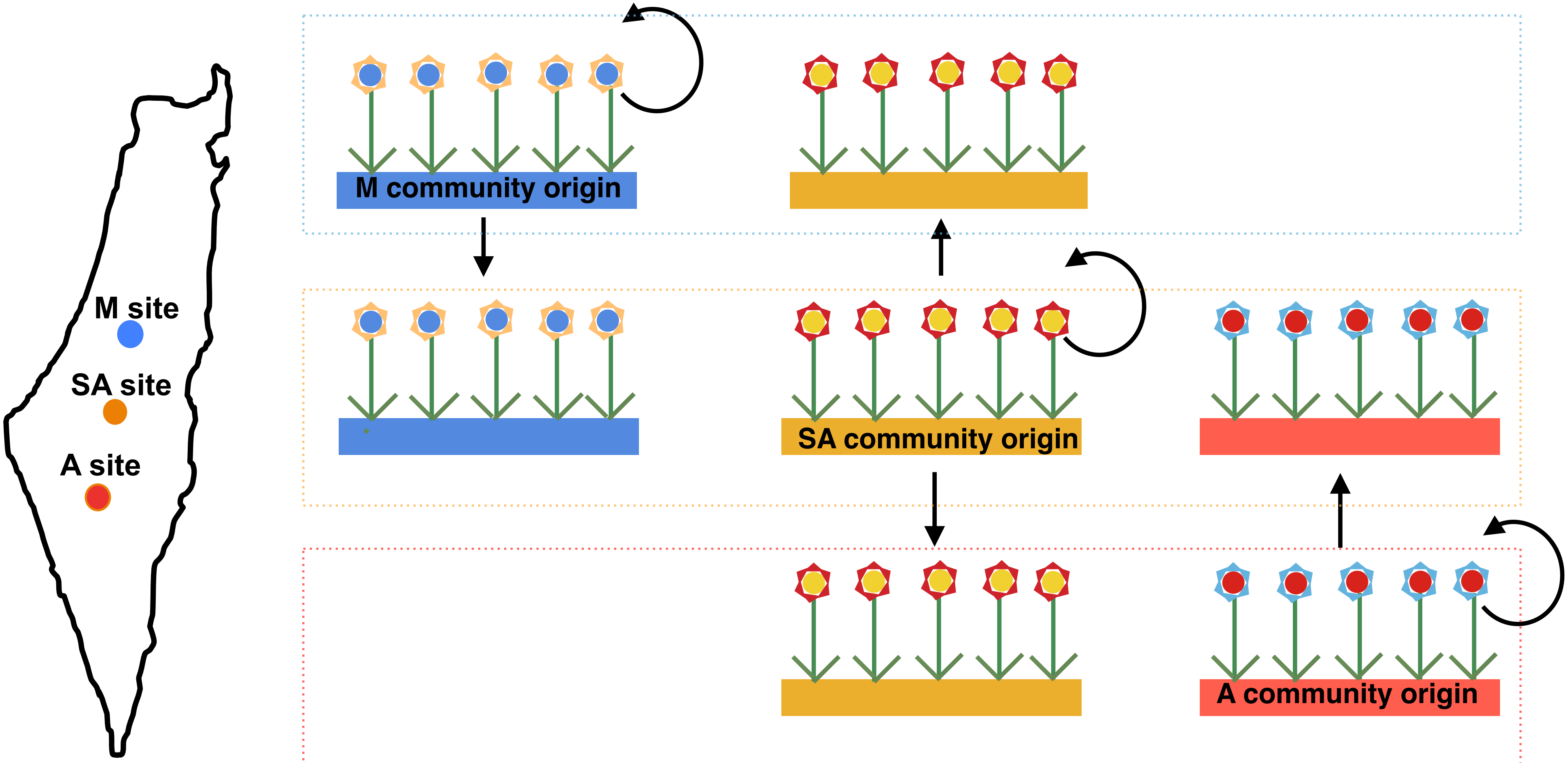
