## Supplementary material for "Whole plant community transplants across climates reveal structural community stability due to large shifts in species assemblage": Species list and mean number of individuals among Climatic Niche Groups

### Appendix 2

**Table A1: List of species and relative Climatic Niche Group (CNG) found in each treatment (i.e. community origin - climate combination). CNG boundaries were set following the classification proposed by Bilton et al (2016). CNG1: 100-230 mm rainfall; CNG2: 230-360 mm; CNG3: 360-490 mm; CNG4: 490-620 mm.**

| Species | Family | Average rainfall (mm/year) | CNG | M-M | SA-M | M-SA | SA-SA |
|---|---|---|---|---|---|---|---|
| *Reichardia tingitana* | Asteraceae | 179.3 | 1 | x | | | x |
| *Stipa capensis* | Poaceae | 221.7 | 1 | x | x | x | x |
| *Adonis dentata* | Ranunculaceae | 227.0 | 1 | | | x | x |
| *Carrichtera annua* | Brassicaceae | 203.2 | 1 | | x | x | x |
| *Plantago coronopus* | Plantaginaceae | 185.5 | 1 | | | | x |
| *Schismus arabicus* | Geraniaceae | 146.9 | 1 | | | x | x |
| *Cuscuta campestris* | Convolvulaceae | 488.1 | 1 | x | x | | |
| *Picris damascena* | Asteraceae | 165.7 | 1 | x | x | x | |
| *Astragalus tribuloides* | Fabaceae | 115.0 | 1 | | x | | |
| *Erucaria rostrata* | Brassicaceae | 168.5 | 1 | | x | | |
| *Filago desertorum* | Brassicaceae | 122.5 | 1 | | | | |
| *Gagea reticultata* | Liliaceae | 143.7 | 1 | | | x | |
| *Pteranthus dichotomus* | Caryophyllaceae | 120.3 | 1 | | x | | |
| *Atractylis cancellata* | Asteraceae | 361.1 | 2 | x | x | x | x |
| *Bromus fasciculatus* | Poaceae | 302.9 | 2 | x | x | x | x |
| *Hippocrepis unisiliquosa* | Fabaceae | 341.3 | 2 | x | x | x | x |
| *Onobrychis crista-galli* | Fabaceae | 279.3 | 2 | x | x | x | x |
| *Trisetaria macrochaeta* | Poaceae | 244.5 | 2 | x | x | | x |
| *Crithopsis delileana* | Poaceae | 302.1 | 2 | | | x | x |
| *Filago contracta* | Brassicaceae | 342.9 | 2 | | x | x | x |
| *Filago palaestina* | Brassicaceae | 344.4 | 2 | | | | x |
| *Helianthemum aegytiacum* | Cistaceae | 360.2 | 2 | x | | | |
| *Helianthemum salicifolium* | Cistaceae | 347.9 | 2 | x | x | x | |
| *Malva parviflora* | Malvaceae | 235.0 | 2 | x | | x | |
| *Astragalus asterias* | Fabaceae | 278.2 | 2 | | x | | |
| *Calendula* sp. | Asteraceae | 349.1 | 2 | | | x | |
| *Hedysarum spinosissimum* | Fabaceae | 347.6 | 2 | | x | | |
| *Aegilops peregrina* | Poaceae | 478.9 | 3 | x | x | x | x |
| *Anagallis arvensis* | Primulaceae | 429.5 | 3 | x | x | x | x |
| *Biscutella didyma* | Brassicaceae | 449.4 | 3 | x | x | x | x |
| *Brachypodium distachyon* | Poaceae | 413.1 | 3 | x | x | x | x |
| *Hymenocarpos circinnatus* | Fabaceae | 443.6 | 3 | x | x | x | x |
| *Plantago afra* | Plantaginaceae | 403.7 | 3 | x | x | x | x |

| | | | | | | | |
|---|---|---|---|---|---|---|---|
| *Plantago cretica* | Plantaginaceae | 468.2 | 3 | x | x | | x |
| *Rhagadiolus stellatus* | Asteraceae | 446.7 | 3 | x | x | x | x |
| *Rostraria cristata* | Poaceae | 469.5 | 3 | x | x | x | x |
| *Sedum pallidum* | Rubiaceae | 438.8 | 3 | x | x | x | x |
| *Medicago truncatula* | Fabaceae | 484.1 | 3 | | x | | x |
| *Medicago tuberculata* | Fabaceae | 484.6 | 3 | | x | | x |
| *Trigonella monspeliaca* | Fabaceae | 438.5 | 3 | | | | x |
| *Vulpia ciliata* | Poaceae | 445.2 | 3 | | | | x |
| *Avena sterilis* | Poaceae | 460.9 | 3 | x | x | x | |
| *Cichorium pumilum* | Asteraceae | 471.9 | 3 | x | | | |
| *Clypeola jonthlaspi* | Brassicaceae | 391.2 | 3 | x | | x | |
| *Convolvulus siculus* | Convolvulaceae | 428.4 | 3 | x | x | x | |
| *Coronilla scorpioides* | Fabaceae | 473.6 | 3 | x | | x | |
| *Crepis sancta* | Asteraceae | 466.1 | 3 | x | | x | |
| *Erodium malacoides* | Geraniaceae | 472.2 | 3 | x | | x | |
| *Galium judaicum* | Rubiaceae | 485.7 | 3 | x | x | x | |
| *Hedypnois rhagadioloides* | Asteraceae | 405.0 | 3 | x | x | x | |
| *Helianthemum stipulatum* | Cistaceae | 373.7 | 3 | x | | | |
| *Linum strictum* | Linaceae | 464.0 | 3 | x | x | x | |
| *Lolium rigidum* | Poaceae | 481.4 | 3 | x | x | x | |
| *Medicago coronata* | Fabaceae | 458.7 | 3 | x | x | | |
| *Onobrychis caput galli* | Fabaceae | 479.5 | 3 | x | x | | |
| *Scorpiurus muricatus* | Fabaceae | 447.8 | 3 | x | x | x | |
| *Torilis tenella* | Apiaceae | 474.0 | 3 | x | x | x | |
| *Urospermum picroides* | Asteraceae | 457.6 | 3 | x | x | x | |
| *Valantia hispida* | Rubiaceae | 448.0 | 3 | x | x | x | |
| *Gynandriris sisyrhinchium* | Iridaceae | 399.4 | 3 | | | x | |
| *Plantago lagopus* | Plantaginaceae | 474.6 | 3 | | x | | |
| *Ranunculus asiaticus* | Ranunculaceae | 437.9 | 3 | | | x | |
| *Cephalaria joppensis* | Dipsacaceae | 574.0 | 4 | x | x | x | x |
| *Hordeum bulbosum* | Poaceae | 540.3 | 4 | x | x | x | x |
| *Medicago rotata* | Fabaceae | 506.4 | 4 | x | x | | x |
| *Sarcopoterium spinosum* | Rosaceae | 511.3 | 4 | x | x | x | x |
| *Trisetaria michelii* | Poaceae | 556.7 | 4 | | | x | x |
| *Althaea hirsuta* | Malvaceae | 602.2 | 4 | x | x | | |
| *Convolvulus pentapetaloides* | Convolvulaceae | 498.3 | 4 | x | x | x | |
| *Crupina crupinastrum* | Asteraceae | 535.6 | 4 | x | x | | |
| *Geranium rotundifolia* | Geraniaceae | 544.9 | 4 | x | | | |
| *Geropogon hybridus* | Asteraceae | 520.8 | 4 | x | | | |
| *Linum corymbosum* | Linaceae | 569.2 | 4 | x | | x | |
| *Linum pubescens* | Linaceae | 557.9 | 4 | x | x | | |
| *Lotus peregrinus* | Fabaceae | 508.5 | 4 | x | x | x | |

| | | | | | | | |
|---|---|---|---|---|---|---|---|
| *Mentha longifolia* | Lamiaceae | 547.4 | 4 | x | | x | |
| *Mercurialis annua* | Euphorbiaceae | 504.1 | 4 | x | | | |
| *Onobrychis squarrosa* | Fabaceae | 509.7 | 4 | x | | | |
| *Scabiosa palaestina* | Dipsacaceae | 515.2 | 4 | x | | | |
| *Scandix iberica* | Apiaceae | 604.9 | 4 | x | | | |
| *Stachys neurocalycina* | Lamiaceae | 576.9 | 4 | x | | | |
| *Theligonum cynocrambe* | Theligonaceae | 515.6 | 4 | x | x | | |
| *Trifolium campestre* | Fabaceae | 502.2 | 4 | x | | x | |
| *Trifolium pilulare* | Fabaceae | 575.2 | 4 | x | x | x | |
| *Trifolium purpureum* | Fabaceae | 526.7 | 4 | x | x | x | |
| *Trifolium scabrum* | Fabaceae | 545.2 | 4 | x | x | x | |
| *Trifolium stellatum* | Fabaceae | 533.0 | 4 | x | x | x | |
| *Tripodion tetraphyllum* | Fabaceae | 536.7 | 4 | x | | | |
| *Vicia palaestina* | Fabaceae | 561.1 | 4 | x | x | | |
| *Bromus japonicus* | Poaceae | 503.8 | 4 | | | x | |
| *Crucianella macrostachya* | Rubiaceae | 494.3 | 4 | | | x | |
| *Nigella ciliaris* | Ranunculaceae | 547.5 | 4 | | x | | |
| *Pallenis spinosa* | Asteraceae | 572.5 | 4 | | x | | |
| *Senecio vernalis* | Asteraceae | 526.9 | 4 | | | x | |
| *Erodium* sp. | Geraniaceae | NA | NA | x | | x | x |
| *Allium* sp. | Liliaceae | NA | NA | | x | x | x |
| *Asphodelus sp.* | Liliaceae | | NA | | x | | |

**Table A2: Mean number ± SE of individuals across Climatic Niche Groups (CNG) emerging in each combination of community origin - climate**

| | | | | |
|---|---|---|---|---|
| **semi-arid community origin** | | | | |
| | **CNG1** | **CNG2** | **CNG3** | **CNG4** |
| **SA (home site)** | 3.55±0.48 | 5.5±0.98 | 5.35±0.73 | 0.30±0.20 |
| **M (away-from-home site)** | 1.57±0.39 | 2.42±0.47 | 8.26±1.41 | 1.94±1.08 |
| **Mediterranean community origin** | | | | |
| | **CNG1** | **CNG2** | **CNG3** | **CNG4** |
| **M (home site)** | 4.0±1.3 | 5.5+0.55 | 39.90+2.21 | 13.55+1.07 |
| **SA (away-from-home site)** | 5.0+0.92 | 5.6+1.06 | 39.55+3.14 | 6.65+0.89 |
